# Mitotic checkpoint protein Mad1 is required for early Nup153 recruitment to chromatin and nuclear envelope integrity

**DOI:** 10.1101/2020.05.20.106567

**Authors:** Ikram Mossaid, Guillaume Chatel, Valérie Martinelli, Marcela Vaz, Birthe Fahrenkrog

## Abstract

The nucleoporin Nup153 is a multifunctional protein and the mitotic checkpoint protein Mad1one of its many binding partners. The functional relevance of their interaction has remained elusive. Here, we have further dissected Nup153’s and Mad1’s interface and functional interplay. By *in situ* proximity ligation assays, we found that the presence of a nuclear envelope (NE) is prerequisite for the Nup153-Mad1 interaction. Time-lapse microscopy revealed that depletion of Mad1 delayed recruitment of Nup153 to anaphase chromatin, which was often accompanied by a prolongation of anaphase. Furthermore, as seen by electron microscopic and three-dimensional structured illumination investigations, Nup153 and Mad1 depletion led to alterations in NE architecture, characterised by a change of the membrane curvature at nuclear pore complexes (NPCs) and an expansion of the spacing between the inner and outer nuclear membranes. Nup153 depletion, but not of Mad1, caused defects in interphase NPC assembly with partial displacement of cytoplasmic nucleoporins and a reduction in NPC density. Together our results suggest that Nup153 has separable roles in NE and NPC formation: in post-mitotic NE reformation in concert with Mad1 and in interphase NPC assembly, independent of Mad1.

**Summary:** The mitotic checkpoint protein is required for Nup153 recruitment to anaphase chromatin and in turn post-mitotic, but not interphase nuclear pore complex assembly.

## Introduction

Nuclear pore complexes (NPCs) are large multi-protein complexes that accomplish all macromolecular exchange across the nuclear envelope (NE). NPCs consist of multiple copies of ~30 different nucleoporins (Nups) that assemble into several biochemically and structurally defined entities: the NPC scaffold, an assembly of distinct ring moieties, the cytoplasmic filaments, and the nuclear basket (Beck and Hurt, 2017; Hoelz et al., 2016; Knockenhauer and Schwartz, 2016; Schwartz, 2016). Major building blocks of the NPC scaffold are the Nup107-160 complex (or Y-complex), which is composed of nine nucleoporins, and the five nucleoporin comprising Nup93 complex. Both, the Nup107-160 and the Nup93 complex are critical for NPC assembly (Boehmer et al., 2003; Hawryluk-Gara et al., 2008; Sachdev et al., 2012; Souquet et al., 2018; Vollmer and Antonin, 2014; Vollmer et al., 2012; Walther et al., 2003). Cytoplasmic filaments and the nuclear basket are predominantly made of phenylalanine-glycine (FG)-repeat containing nucleoporins, which are of particular importance for nucleocytoplasmic transport (Lim et al., 2008; Patel et al., 2007; Terry and Wente, 2009). One such FG-repeat nucleoporin is Nup153, a constituent of the nuclear basket (Fahrenkrog et al., 2002; Pante et al., 2000; Walther et al., 2001). Nup153 is a multi-functional protein with tasks that go well beyond nucleocytoplasmic transport (Ball and Ullman, 2005; Duheron et al., 2017; Lemaitre et al., 2012; Lussi et al., 2010; Mackay et al., 2009; Mackay et al., 2017; Moudry et al., 2012; Nanni et al., 2016; Prunuske et al., 2006; Toda et al., 2017; Vaquerizas et al., 2010; Zhou and Pante, 2010). In this regards, Nup153 is known to play a role in both, post-mitotic and interphase NPC assembly.

Post-mitotic NPC assembly is initiated as early as anaphase, and the first nucleoporins that accumulate on chromatin are members of the Nup107-160 complex followed by a small fraction of Nup153 and its nuclear basket partner Nup50 (Anderson et al., 2009; Dultz et al., 2008; Schwartz et al., 2015; Vagnarelli and Earnshaw, 2012). Nup107-160 recruitment is mediated by ELYS/Mel-28, which associates to chromatin via its AT-hook DNA binding motif (Fernandez and Piano, 2006; Franz et al., 2007; Lau et al., 2009; Rasala et al., 2008; Walther et al., 2003). Recruiting Nup153 onto chromatin occurs independent of ELYS and may substitute for ELYS to recruit the Nup107-160 complex (Schwartz et al., 2015). Nup153 is capable of recruiting a large number of nucleoporins from basically all NPC substructures (Bilir et al., 2019; Dultz et al., 2008; Schwartz et al., 2015; Walther et al., 2001), indicating that it is able to seed the formation of NPCs on chromatin (Schwartz et al., 2015). Important for targeting Nup153 on anaphase chromatin is a complex between Repo-Man and importin ß (Vagnarelli and Earnshaw, 2012; Vagnarelli et al., 2011). Lack of Repo-Man impairs importin ß and subsequent recruitment of Nup153 to the anaphase chromatin. Whether Repo-Man docks to the chromatin directly or indirectly has remained unclear (Vagnarelli and Earnshaw, 2012).

Interphase NPC assembly occurs into a closed NE as cells progress through interphase (Maul et al., 1971; Otsuka and Ellenberg, 2018) and it requires the Nup107-160 complex (D’Angelo et al., 2006; Doucet and Hetzer, 2010; Doucet et al., 2010; Harel et al., 2003; Walther et al., 2003). Similarly, the transmembrane nucleoporin POM 121 and Nup153 are necessary (Talamas and Hetzer, 2011; Vollmer et al., 2015), but not so ELYS (Doucet et al., 2010). Interphase NPC assembly requires an insertion into the nuclear membrane and fusion of the outer nuclear membrane (ONM) and the inner nuclear membrane (INM). Nup153 can interact with the INM via an N-terminal amphipathic helix and Nup153 insertion into the INM facilitates the recruitment of the Nup107-160 complex to these NPC assembly sites (Vollmer et al., 2015). Likewise, Nup1, the yeast paralog of Nup153, induces membrane curvature by amphipathic helix insertion into the lipid bilayer (Meszaros et al., 2015). The interaction of Nup153 with the INM is regulated by its nuclear import receptor transportin and by the small GTPase RanGTP (Vollmer et al., 2015).

Mad1 is a key component of the mitotic checkpoint (or spindle assembly checkpoint (SAC)), which delays anaphase onset until all chromosomes have been properly attached to the mitotic spindle. Mad1 localises to NPCs in interphase (Campbell et al., 2001; Iouk et al., 2002; Lussi et al., 2010), where it binds the two nuclear basket nucleoporins Tpr and Nup153 (Ding et al., 2012; Lee et al., 2008). The association of Mad1 with Tpr plays an important role in mitotic checkpoint regulation (Cunha-Silva et al., 2020; Lee et al., 2008; Lopez-Soop et al., 2017; Rajanala et al., 2014; Rodriguez-Bravo et al., 2014; Schweizer et al., 2013), whereas the functional significance of the Mad1-Nup153 complex has remained largely elusive. Here we provide evidence that Mad1 is required for seeding Nup153 on anaphase chromatin and post-mitotic NPC assembly.

## Results

### Mad1 exhibits two independent binding sites for Nup153

We have previously shown that Nup153 and Mad1 are directly interacting, both *in vitro* and *in situ*, and that the N-terminal domain of Nup153 establishes the binding to Mad1 (Lussi et al., 2010). To deepen the analysis of the Nup153-Mad1 interface and to identify the region of Mad1 that binds Nup153, we carried out GST pull-down assays. To do so, we expressed distinct Mad1 fragments fused to a FLAG tag (Fig. 1A) and the N-terminal part of Nup153 fused to GST (residues 2-610; (Lussi et al., 2010); GST-153N) in *E. coli*. GST alone was used as negative control. As shown in Fig. 1B, full-length FLAG-Mad1 (Mad1) and the N-terminal domain of Mad1 (residues 1-596; N596) bound to GST-Nup153, but not to GST. Mad1’s C-terminal domain (CTD, residues 597-718; C597-718) showed no interaction with GST-Nup153. A further dissection of Mad1’s N-terminal domain revealed that (i) the Mad2-interacting motif (MIM) of Mad1 (N539) is dispensable for binding Nup153 (Fig. 1C); and that (ii) residues 121-240 (N121-240) are sufficient for Nup153 binding (Fig. 1D). Consequently Mad1 residues 1 to 240 (N240; Fig. 1C) bound Nup153, in contrast to fragments comprising residues 1-120 (N120) and residues 241-539 (N241-539; Fig. 1D). These findings are in good agreement with a previozs study that showed that residues 1-274 are necessary and sufficient for the localisation of Mad1 at NPCs (Rodriguez-Bravo et al., 2014).

**Figure 1:**
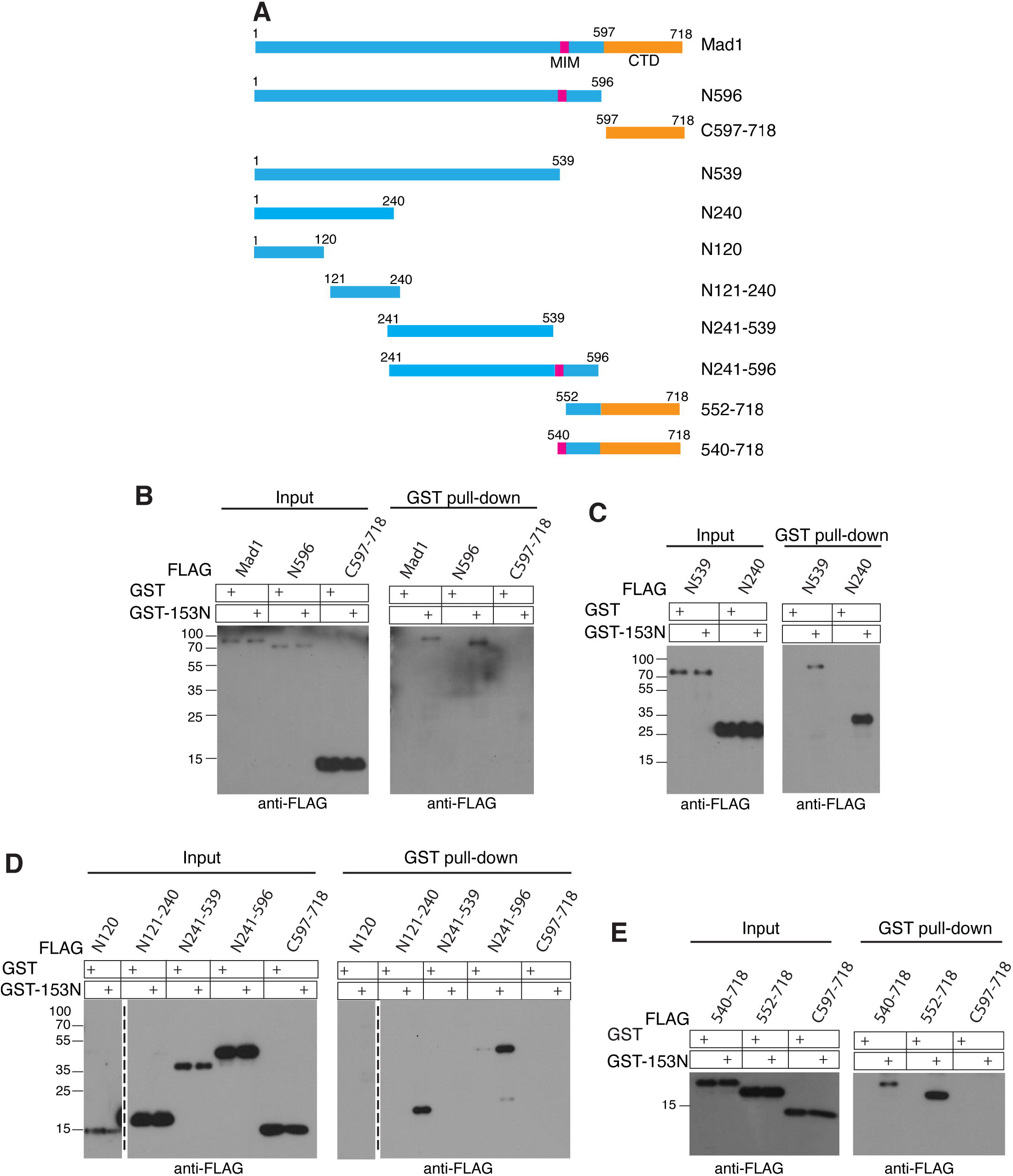
Mad1 owns two independent binding sites for Nup153. **(A)** Schematic representation of Mad1 fragments used for binding assays. (**B**) GST pulldown-assays were performed using recombinant GST-Nup153-N and GST alone. Recombinant FLAG-tagged Mad1 fragments comprised full-length Mad1, a fragment of N-terminal domain (residues 1-596; N596) and a fragment of the C-terminal domain (residues 597-718; C597-718). Input and bound fractions (GST pull-down) were analysed by immunoblotting using anti-FLAG antibodies. Analysis of the interaction between Nup153 and shorter N-terminal fragments of Mad1 (**C**) comprising residues 1-539 (N539) and residues 1-240 (N240), (**D**) residues 1-240 (N120), residues 121-240 (N121-240), residues 241-539 (N241-539), and residues 241-596 (N241-596). Nup153 interacts residues 1-240, the N121-240, and the N241-596 fragments of Mad1. (**E**) GST pull-down using Mad1 fragments spanning residues 540-718 (540-718), residues 552 to 718 (552-718; lacking the MIM domain), and C597-718. Mad1 residues 552-596 mediate the interaction with Nup153.

We further identified a second Nup153 binding site in the N terminus of Mad1: residues 241-596, 540-718, and 552-718 of Mad1 did bind Nup153-N (Fig. 1D and E), but not so residues 597-718 (Fig. 1B, E). We therefore concluded that the N-terminal domain of Mad1 comprises two binding sites for Nup153: one involving residues 121 to 240 and the second residues 552 to 596.

### Nup153 and Mad1 interact exclusively in the presence of the NE

While Nup153 and Mad1 co-localise at NPCs during interphase (Lussi et al., 2010), they adopt different locations during mitosis: Nup153 is found dispersed in the mitotic cytoplasm (Dultz et al., 2008; Mackay et al., 2009), whereas Mad1 is localised to unattached kinetochores during prophase and prometaphase to fulfil its SAC function (Chen et al., 1998). To further characterise the interaction between Nup153 and Mad1 in space and time in cells, we next performed *in situ* proximity ligation assays (PLA) in HeLa cells (see Materials and Methods; (Soderberg et al., 2006)). Abundant PLA signals at the edge of the nucleus (visualised by DAPI staining) were detected in interphase cells (Fig. 2A). PLA signals were diminished upon depletion of Nup153 or Mad1 (Fig. 2A, B). Depletion of Tpr, another nucleoporin known to bind Mad1 (Lee et al., 2008), had no impact on the Nup153-Mad1 PLA signals (Fig. 2A, B), suggesting that the Nup153 and Mad1 interaction is independent of Tpr. PLA results were validated by different control pairs (Fig. S1A, B). Knock-down efficiency for the distinct siRNAs was determined by Western blot (Fig. S1C, D).

**Figure 2:**
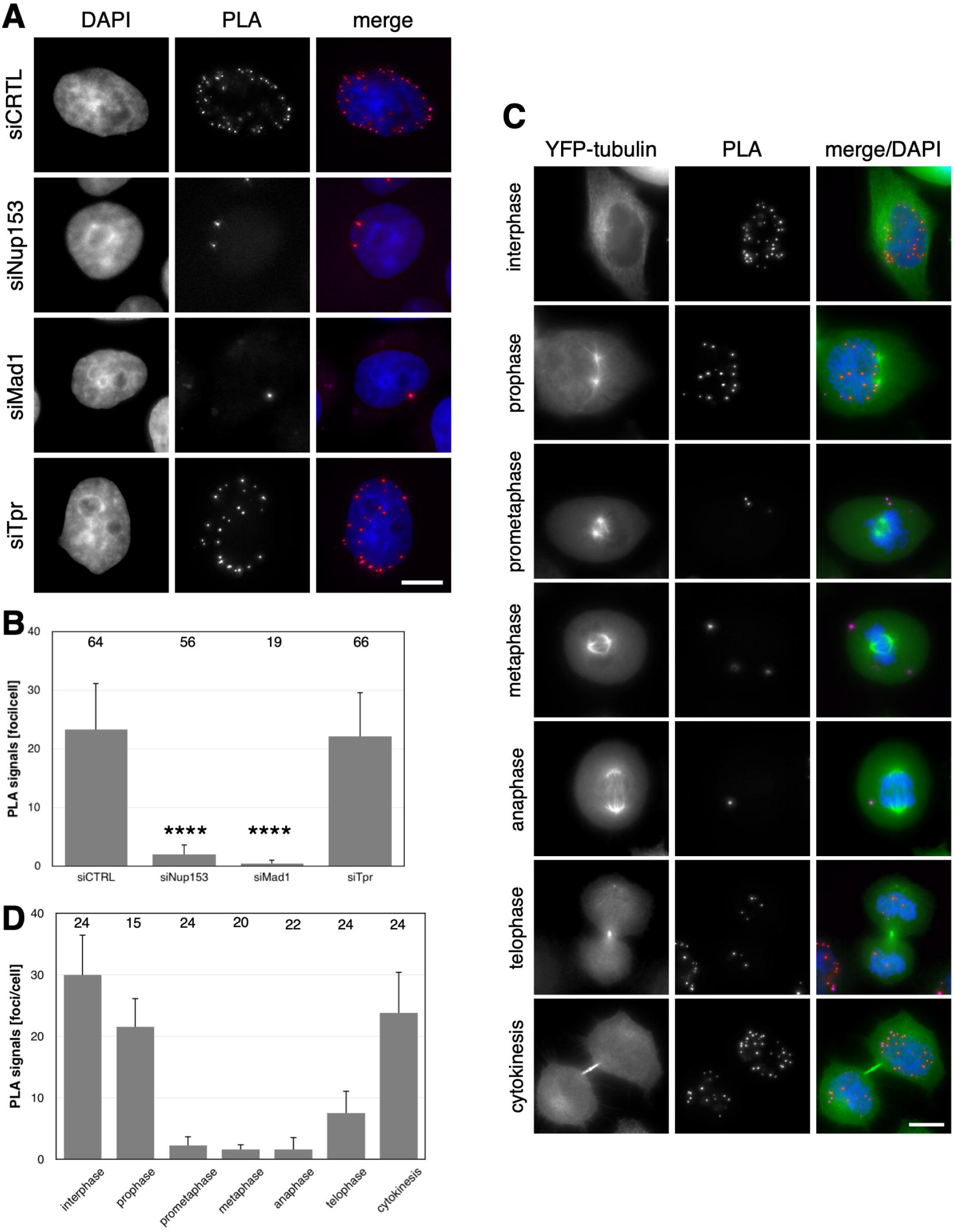
Proximity ligation assays (PLA) revealed a nuclear envelope-dependent association between Nup153 and Mad1. (**A**) HeLa cells were treated with the indicated siRNAs for 48 h (siTpr for 72 h) and labelled with primary anti-Nup153 and anti-Mad1 antibodies followed by secondary oligonucleotide-linked probes. DNA (blue) was visualised by DAPI. Shown are representative epifluorescence images. Scale bar, 10 μm. (**B**) Quantification of Nup153-Mad1 PLA foci per cell after treatment with the indicated siRNAs. Total numbers of analysed cells per condition are indicated at the top of the graph. Values are mean ± SD.**** p<0.0001; t-test, two-tailed. (**C**) HeLa cells stably expressing YFP-tubulin subjected to *in situ* PLA at different cell cycle states using anti-Nup153 and anti-Mad1 antibodies. PLA association (red) between Nup153 and Mad1 primarily from telophase to prophase, i.e. in the presence of a nuclear envelope. Shown are representative epifluorescence images. DNA (blue) was visualised by DAPI. Scale bar, 10 μm. (**D**) Quantification of Nup153-Mad1 PLA foci for the distinct cell cycle states. Total numbers of analysed cells per condition are indicated at the top.

Having confirmed the interaction between Nup153 and Mad1 at the NE, we next asked whether Nup153 and Mad1 show some association during mitosis, which may have previously escaped detection by conventional immunofluorescence microscopy. We performed PLA assays in HeLa cells stably expressing YFP-tubulin to precisely monitor the cell cycle state. As shown in Fig. 2C, PLA signals for Nup153 and Mad1 were only visible in the presence of the NE. Quantification of the PLA signals for each cell cycle state revealed that PLA signals were established in telophase when the NE is forming, increased during cytokinesis, when NE formation is completed, remained during interphase, and declined in prophase, when the NE begins to disassemble (Fig. 2D). Consistent with these data we observed no significant PLA between Nup153 and Mad1 during prometaphase in normal HeLa cells (Fig. S1B). Additionally, no PLA between Nup153 and the outer kinetochore protein Hec1 (Wei et al., 2005) was observed during prometaphase, in contrast to Mad1 and Mad2 (Fig. S1B). Together these data confirm the absence of Nup153 from mitotic structures (Mackay et al., 2009) and they suggest a dissociation of the Nup153-Mad1 complex at the beginning of the mitosis when the NE disassembles.

### Mad1 is required for Nup153 recruitment to the reforming NE

We previously reported that overexpression of Nup153 caused a SAC override, whereas depletion of Nup153 had no obvious impact on SAC function (Lussi et al., 2010). This observation is confirmed by decreasing phospho-histone H3 levels, timely cyclin B1 and securin degradation (Fig. S2A), as well as timely Cdc20 dissociation from Mad2 in Nup153-depleted cells after release from nocodazole arrest (Fig. S2B). Because of a missing apparent effect of Nup153-depletion on Mad1/SAC function, we next assessed whether Mad1-depletion impacts Nup153. By immunofluorescence experiments, we found that the recruitment of Nup153 to chromatin in anaphase was compromised in the absence of Mad1 (Fig. 3A). On the contrary, Nup153’s localisation in any other cell cycle phase was indistinguishable between control and Mad1-depleted cells (Fig. 3A). The effect of Mad1 depletion on Nup153 recruitment to anaphase chromatin appeared specific: in the absence of Mad1 neither anaphase recruitment of importin β to chromatin (Fig. S3A; see also (Vagnarelli et al., 2011)) was altered, nor Tpr recruitment in late telophase (Fig. S3B).

**Figure 3:**
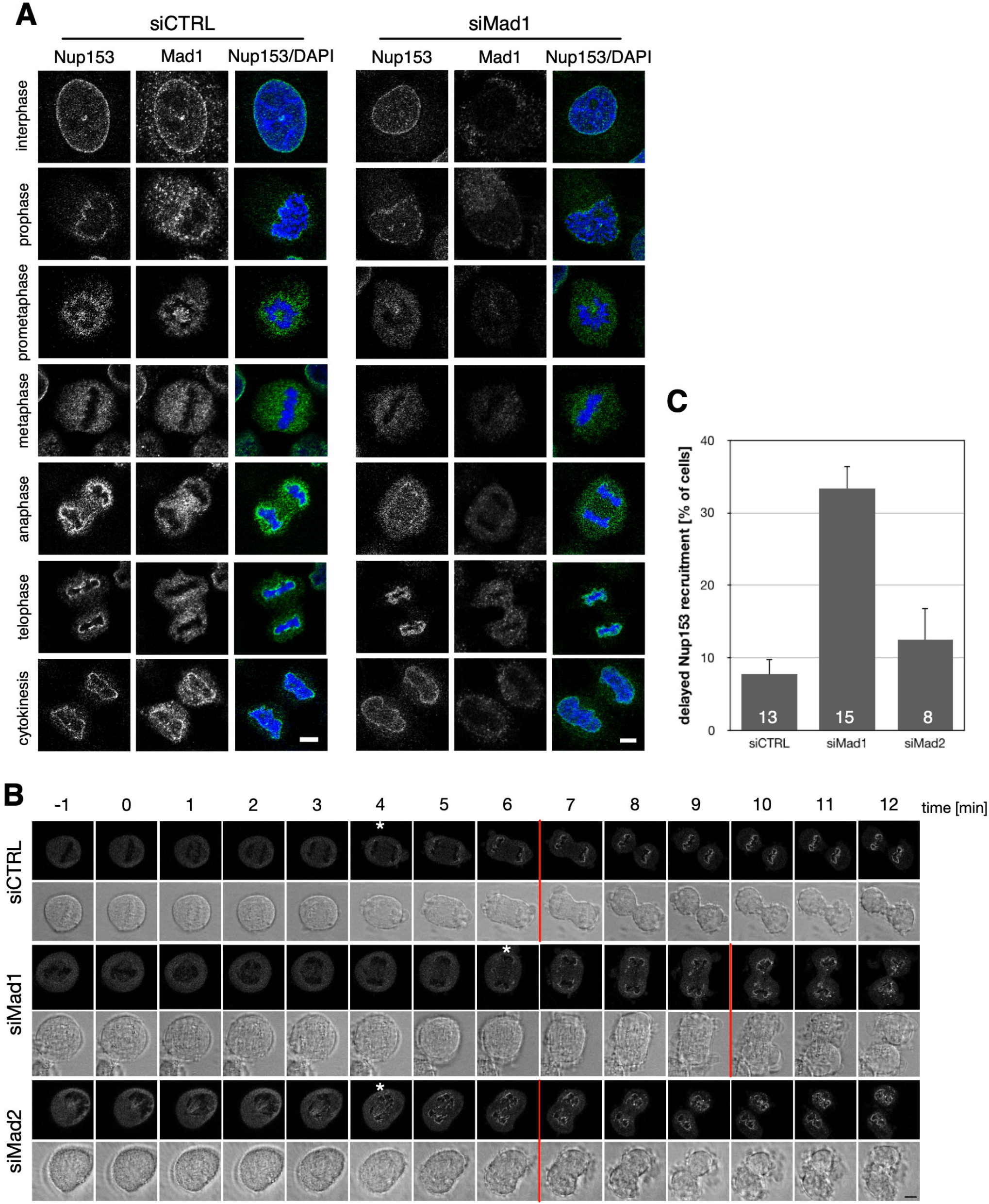
Mad1 depletion delays recruitment to chromatin in anaphase. (**A**) HeLa cells were transiently transfected with the indicated siRNAs, labelled with anti-Nup153 and anti-Mad1 antibodies and analysed by confocal laser-scanning microscopy. In Mad1-depleted cells, Nup153’s recruitment to chromatin is delayed. DNA was visualised by DAPI. Scale bars, 5 μm. (**B**) HeLa T-Rex cells expressing GFP-Nup153 were transiently transfected with the indicated siRNAs and subjected to live imaging 48 hours after transfection. Imaging started two minutes (−1) before anaphase onset (1) using confocal laser-scanning microscopy. Images were taken every minute from the last time point before anaphase onset. White arrows indicate the first evidence of Nup153 recruitment to the condensed chromatin, red bars the onset of telophase. Shown are differential interference contrast and confocal images. Scale bar, 5 μm. (**C**) Quantification of cells exhibiting a delay of Nup153 recruitment to chromatin. Total number of analysed cells per condition is indicated at the bottom of each bar.

Time lapse imaging in HeLa T-Rex cells conditionally expressing GFP-Nup153 (Duheron et al., 2014) confirmed the necessity of Mad1 for Nup153 recruitment to anaphase chromatin. Cells were imaged every minute from anaphase to cytokinesis (representative images are shown in Fig. 3B). While Nup153 appeared on chromatin within 4 minutes (white asterisk) after anaphase onset in control cells, it took about 6 minutes in Mad1-depleted cells. This delay in Nup153 recruitment often coincided with an elongation of anaphase (Fig. 3B, red line) and was observed in about 33% of Mad1-depleted cells, but only in about 7% of control cells (Fig. 3C). Mad2 depletion on the contrary had no effect on Nup153 recruitment to chromatin in anaphase and the duration of anaphase (Fig. 3B,C). Similarly, Mad1 depletion had no effect on the recruitment of GFP-Nup98 to the reforming NE, i.e. 1-2 minutes after the end of anaphase (Fig. S3C), indicating that the effect of Mad1 depletion on Nup153 is specific.

### Nup153 and Mad1 are required for nuclear envelope integrity

We next performed electron microscopy analysis of Nup153- and Mad1-depleted HeLa cells. Strikingly, the space between outer and inner nuclear membrane, i.e. the perinuclear space (PNS), was significantly increased in Nup153- and Mad1-depleted cells as compared to control cells (Fig. 4A,B). We furthermore noticed that the membrane curvature at NPCs was reduced in Nup153- and Mad1-depleted cells (Fig 4B, black arrow). The PNS increased from about 40-50 nm in control cells to about 80-100 nm in Nup153- and Mad1-depleted cells (Fig. 4C). PNS spacing and membrane curvature remained unaffected by the depletion of Nup50 and Tpr, two other well-known Nup153 binding partners at the nuclear basket (Duheron et al., 2014), as well as by the depletion of the three cytoplasmic located nucleoporins Nup358, Nup214, and Nup88 (Fig. S4A). We confirmed the dilation of the PNS upon the respective depletion of Nup153 and Mad1 by three-dimensional structured illumination microscopy (3D-SIM). The ONM was visualised by anti-Nesprin-2 antibodies (Fig. 4D, green) and the INM by anti-lamin A/C antibodies (Fig. 4D, red). While both signals appeared in the same focal plane in control cells, green and red signals were partially separated in the respective absence of Nup153 and Mad1, indicating a spacing of the two membranes of more than 100 nm, i.e. the optical resolution of SIM. This increase in spacing between the ONM and the INM, however, appeared to have no effect on the localisation of nuclear membrane proteins, as indicated by normal NE localisation of Sun1 (ONM) and Nesprin2 (INM; Fig. S4B).

**Figure 4:**
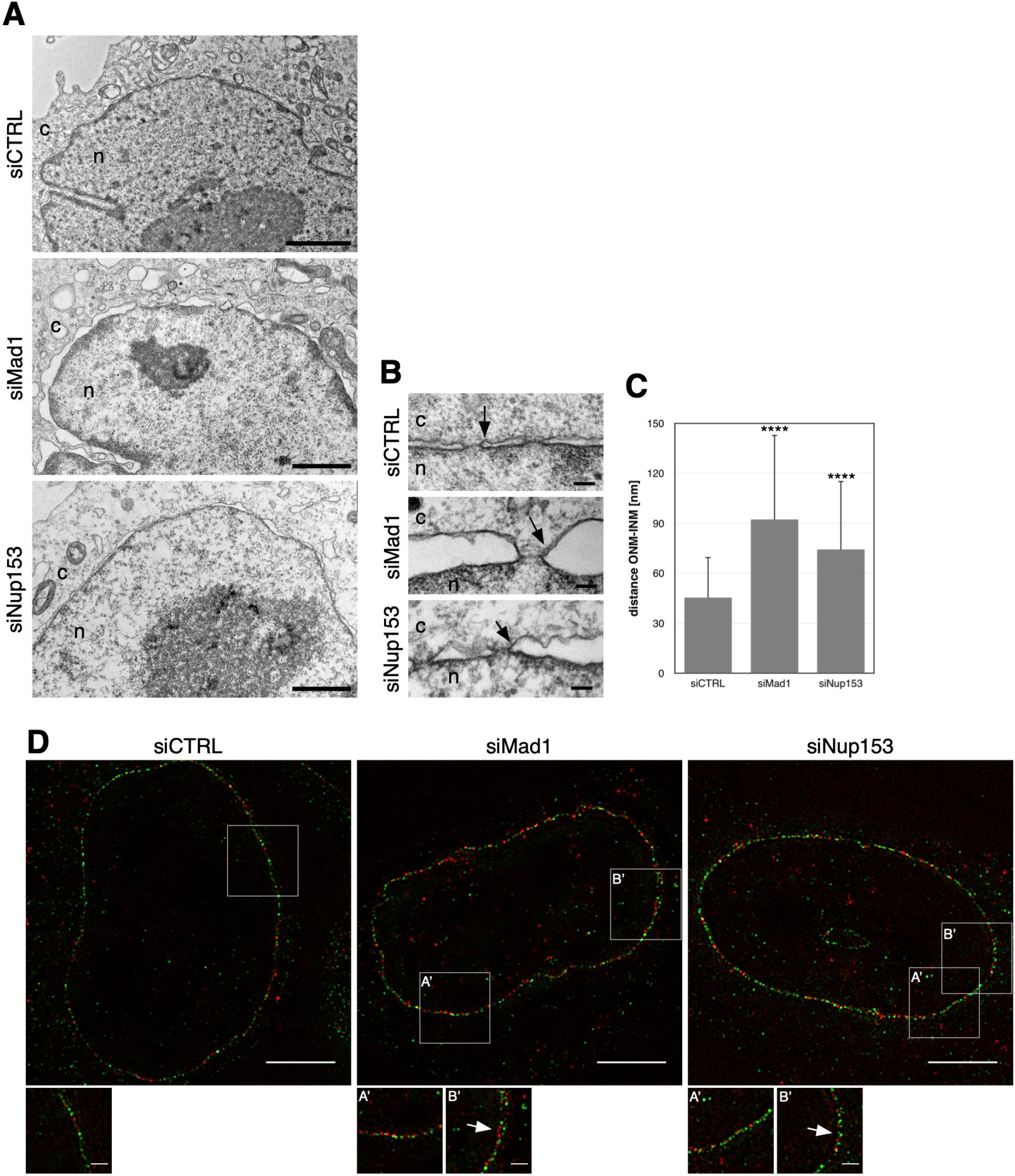
Nup153 and Mad1 depletion induces nuclear envelope anomalies. (**A**) HeLa cells were treated with the indicated siRNAs for 48 h and prepared for thin-sectioning electron microscopy. Shown are cross-sections along the NE. (**B**) NE cross section at higher magnification. The perinuclear space is enlarged in Mad1- and Nup153-depleted cells and membrane curvature is altered (black arrows). Scale bars, 500 nm (**A**), 100 nm (**B**). n, nucleoplasm; c, cytoplasm. **(C)** Quantification of the spacing between ONM and INM of Nup153- and Mad1-depleted cells. 10 nuclei were analysed per condition and NE width was measure at 10 different points per nucleus. **** p< 0.0001; t-test, two-tailed, Mann-Whitney. **(D)** HeLa cells transiently transfected with Nup153 and Mad1 siRNAs, stained with antibodies against Nesprin-2 (green) and lamin A/C (red), and analysed by 3D-SIM. Zoom-in images of the highlight area show the increased distance between Nesprin-2 and lamin A/C (white arrows) upon depletion of Nup153 and Mad1. Scale bars, 5 μm; insets 1 μm.

### Nup153, but not Mad1, is required for NPC integrity

In yeast, Nup1, the functional homolog of Nup153, is known to bind the NE and to regulate membrane curvature due to an amphipathic helix in its N-terminal part (Meszaros et al., 2015). This helix, which is able of interacting with lipids, is implicated in the membrane remodelling process and the deletion of this N-terminal part of Nup1 provoked a reduction of the membrane curvature. This reduced membrane curvature coincided with a partial mis-localisation of the cytoplasmic nucleoporins Nup159 and Nup82 (Meszaros et al., 2015). We wondered if the reduction of the membrane curvature observed in Nup153-depleted HeLa cells would alter cytoplasmic nucleoporins in mammalian cells as well. We therefore monitored the localisation of the cytoplasmic ring nucleoporins Nup214 (ortholog of the yeast Nup159) and Nup88 (ortholog of yeast Nup82) as well as the mammalian cytoplasmic filament protein Nup358 by immunofluorescence microscopy. As shown in Fig. 5A, Nup153 depletion led to a partial displacement of Nup214, Nup88, as well as Nup358 from the NPCs and to their accumulation in cytoplasmic foci. Mad1 depletion on the contrary did not alter the localisation of the three cytoplasmic nucleoporins (Fig. 5B).

**Figure 5:**
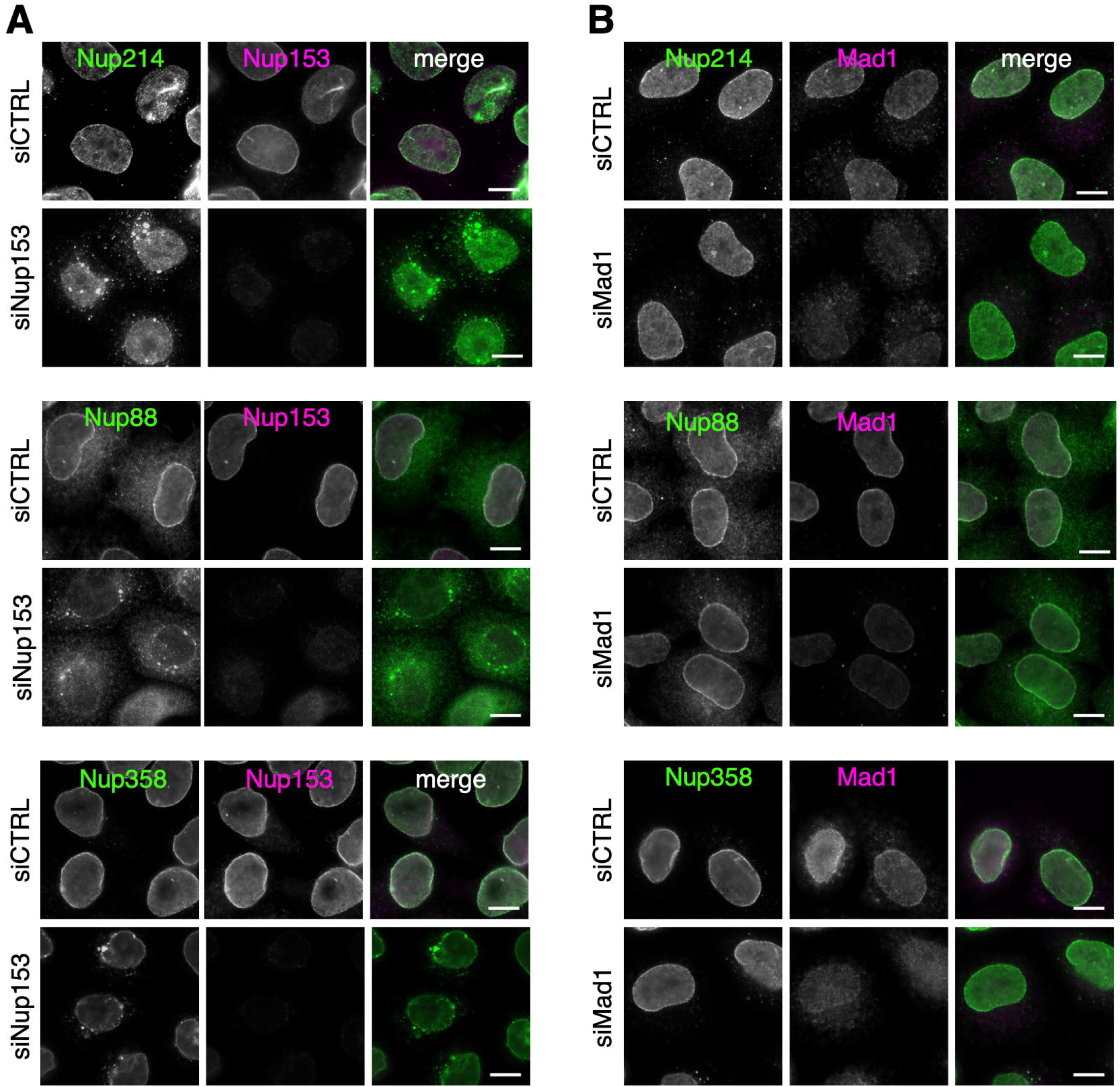
Partial displacement of cytoplasmic nucleoporins from NPCs upon Nup153 depletion. **(A)** siRNA-mediated depletion of Nup153 caused a partial displacement of Nup88, Nup214, and Nup358 to foci in the cytoplasm, in contrast to **(B)** Mad1 depletion. HeLa cells were treated with the indicated siRNAs and labelled with anti-Nup214, anti-Nup88, and anti-Nup358 antibodies (green) and co-stained with anti-Mad1 antibodies (magenta). Scale bars,10 μm.

Furthermore, we extended this analysis to mouse 3T3 fibroblast cells to investigate whether the effect of the depletion of Nup153 is conserved in mammalian cells. In a manner similar to HeLa cells, we detected a mis-localisation of Nup214 and Nup358 in Nup153-depleted 3T3 cells (Fig. 6). To confirm that the effect is specifically caused by Nup153 depletion, we next carried out rescue experiments by expressing human GFP-Nup153 in control and Nup153-depleted 3T3 cells. Human and mouse Nup153 share 83% identity on protein level (www.ncbi.nlm.nih.gov/homologene). As shown in Fig. 6A, Nup214 and Nup358 accumulate in cytoplasmic foci in control and Nup153-depleted 3T3 cells expressing GFP, while expression of human GFP-Nup153 restored Nup214’s and Nup358’s localisation to NPCs in Nup153-depleted 3T3 cells (Fig. 6B). Depletion of endogenous mouse Nup153 and expression of human GFP-Nup153 was confirmed by Western blot analysis (Fig. 6C). Altogether, our data suggest that Nup153 may play an essential, conserved function in NPC assembly in mammalian cells, similar to its function in yeast.

**Figure 6:**
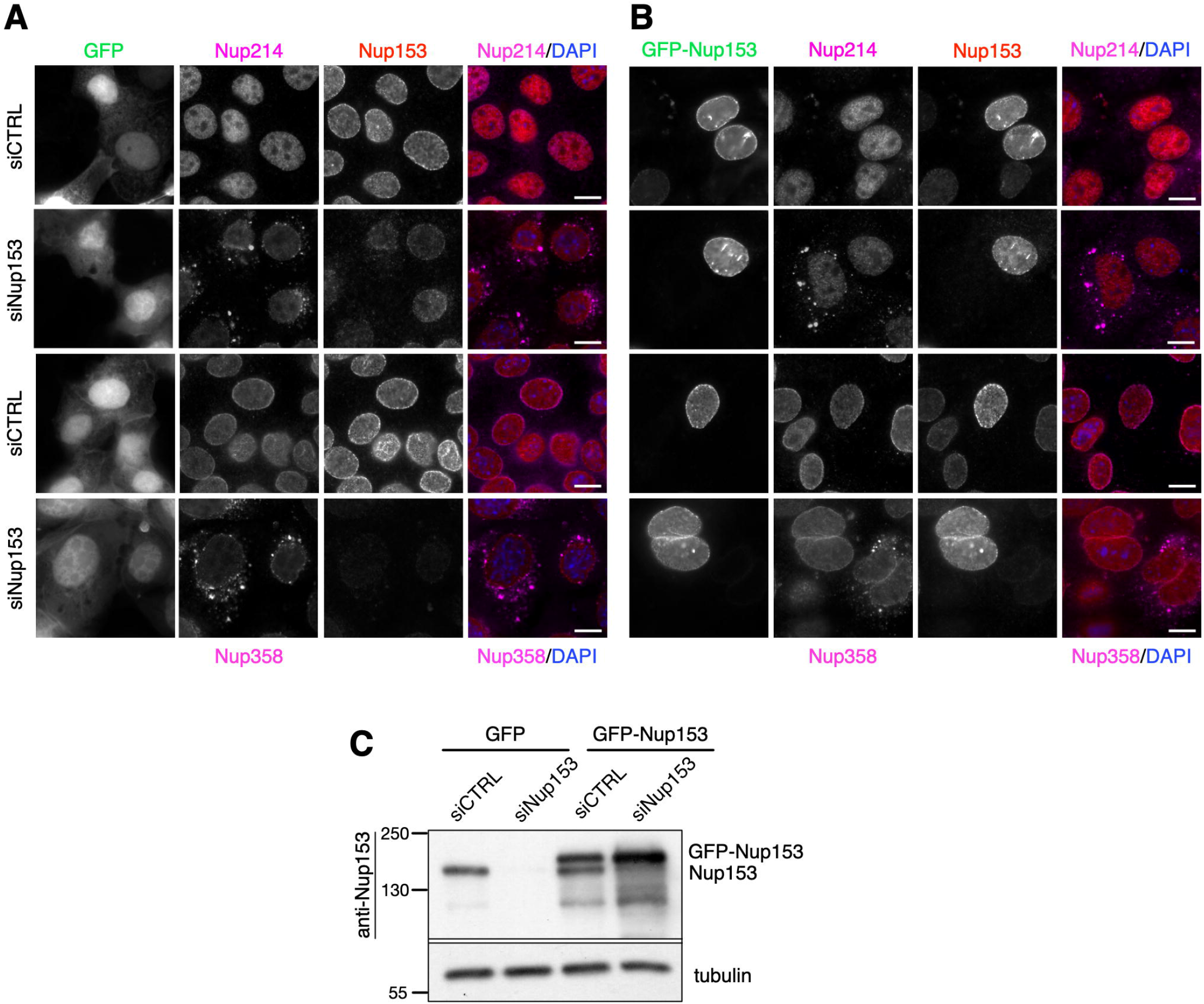
GFP-Nup153 restores the localisation of cytoplasmic nucleoporins in Nup153-depleted cells. Mouse 3T3 cells were treated with control and Nup153 siRNAs for 48 h, transfected with GFP and GFP-Nup153 for 24 h, fixed and stained with (**A**) anti-Nup214 and (**B**) anti-Nup358 antibodies (magenta) and co-stained with anti-Nup153 antibodies (red). The localisation of Nup214 and Nup358 is restored in Nup153-depleted cells expressing RNAi resistant GFP-Nup153. Shown are epifluorescence images. Scale bars, 10 μm. (**C**) Western blot analysis showing the expression of Nup153 in total cell protein lysates from control or Nup153-depleted cells expressing GFP or GFP-Nup153. Anti-α-tubulin antibodies were employed as loading control.

### Nup153 depletion affects NPC assembly in interphase cells

Having seen that depletion of Nup153 leads to a partial mis-localisation of the cytoplasmic nucleoporins, we next investigated whether this defect results from a defect in interphase NPC assembly or at the end of the mitosis. Interphase NPC assembly occurs after DNA replication in S phase. We therefore arrested cells before entry into mitosis in the G2/M boundary by treatment with the CDK1 inhibitor RO-3306. Non-synchronised and arrested cells were stained with NPC-specific mAb414 antibodies, which recognize nucleoporins containing FG-repeats. As shown in Fig. 7A and quantified in Fig. 7B, NPC density was significantly reduced in RO-3306-arrested, Nup153-depleted cells. Mad1 depletion on the contrary did not affect NPC density, suggesting that Nup153, but not Mad1, is required for interphase NPC assembly.

**Figure 7:**
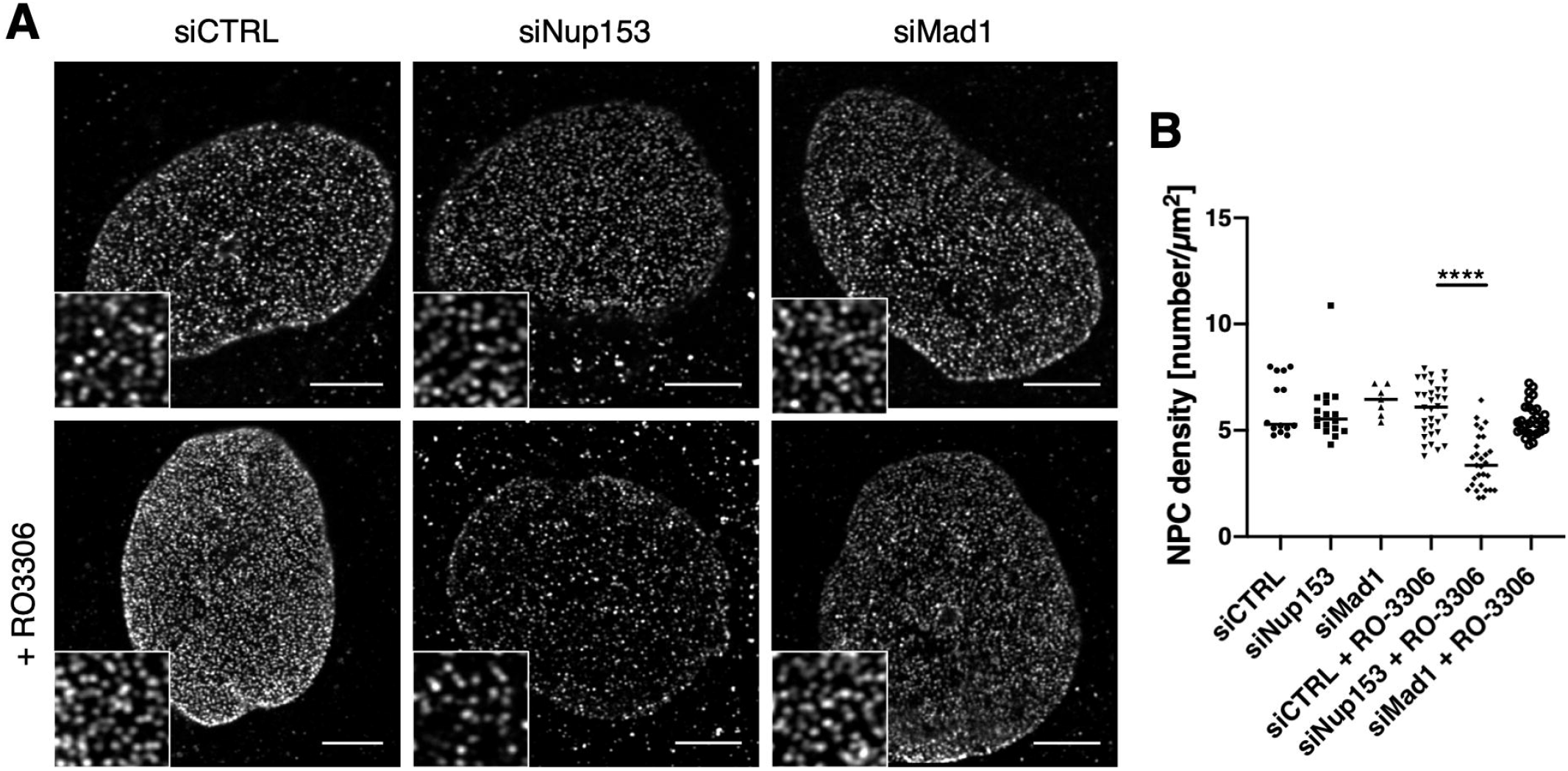
NPC density is reduced in Nup153-depleted cells. **(A)** HeLa cells were transfected with the indicated siRNAs and 48 hours post transfection treated with RO-3306 for 18 h to induce a G2/M arrest. NPCs were visualised by mAb414 antibodies. Shown are representative confocal images in combination with a Zeiss Airyscan detector. Scale bars, 5 μm. **(B)** Quantification of NPC numbers on the nuclear surface using the Fiji/ImageJ cell counter plugin. One-way Anova test, ****p <0.0001, **p <0.01, p <0.05.

## Discussion

We previously identified the spindle checkpoint protein Mad1 as binding partner of Nup153 (Lussi et al., 2010). In the present study, we confirm this interaction and establish a role for Mad1 in Nup153 recruitment to chromatin in anaphase and in post-mitotic NPC assembly.

Our previous study has shown that the N-terminal domain of Nup153 binds Mad1 (Lussi et al., 2010). Mad1 comprises two independent binding sites for this N-terminal region of Nup153: the first region involves residues 121-240, the second region residues 552-596 (Fig. 1). Residues 121-240 are overlapping with the previously described NPC targeting domain of Mad1 (residues 1 to 274), which mediates its interaction with Tpr (Rodriguez-Bravo et al., 2014). Residues 552-596 are downstream of the Mad2-interacting motif (MIM; Fig. 1), in agreement with the previous observation that Mad2 is not co-purifying with the Nup153-Mad1 complex (Lussi et al., 2010). Along this line and consistent with previous studies (Lussi et al., 2010; Mackay et al., 2009; Mackay et al., 2010) depletion of Nup153 had no obvious impact on the SAC (Fig. S2), in contrast to the Tpr-Mad1complex (Lee et al., 2008; Schweizer et al., 2013). Furthermore, our PLA experiments revealed that Nup153 and Mad1 bind each other exclusively in the presence of the NE, i.e. from late telophase to early prophase (Fig. 2C). We confirmed that Nup153 and Mad1 are not co-localising to kinetochores during prometaphase (Lussi et al., 2010; Mackay et al., 2010), therefore we consider it highly unlikely that Nup153 and Mad1 act together in SAC control.

Surprisingly, Mad1 depletion led to delayed recruitment of Nup153 to chromatin in anaphase, coinciding with a prolonged anaphase (Fig. 3B). At this stage we do not know whether Mad1 directly targets Nup153 to chromatin or whether it bridges another factor. Known to be important for targeting Nup153 on anaphase chromatin is a complex between Repo-Man and importin ß (Vagnarelli and Earnshaw, 2012; Vagnarelli et al., 2011). Lack of Repo-Man impairs importin ß and subsequent recruitment of Nup153 to the anaphase chromatin. Whether Repo-Man docks to the chromatin directly or indirectly has remained unclear (Vagnarelli and Earnshaw, 2012). Repo-Man also targets protein phosphatase 1γ (PP1γ) to chromatin in anaphase and regulates chromosome remodelling during the late stages of mitosis. When PP1γ is absent, anaphase is extended (Axton et al., 1990; Chen et al., 2007), similarly to what we observed in Mad1-depleted cells (Fig. 3B). Delayed recruitment of Nup153 in Mad1-depleted cells might therefore arise from a perturbed Repo-Man-PP1γ localisation or function. Although we did not observe a mis-localisation of importin β in Mad1-depleted cells (Fig. S3B), a Repo-Man-mediated effect of Mad1 on Nup153 cannot be completely ruled out. Alterations in such short-lived dynamic processes as recruitment to chromatin during anaphase might be missed in fixed samples and will require in-depth time-lapse imaging to monitor localisation of Repo-Man and PP1γ on chromatin during anaphase in Mad1-depleted cells.

What is the consequence of failed recruitment of Nup153 to anaphase chromatin: at first sight relatively little. While Nup153 acts as seed for NPC assembly (Schwartz et al., 2015), it is not the only seeding nucleoporin. Nup133, ELYS, and Nup50 are equally capable of seeding NPCs (Schwartz et al., 2015), indicating a highly secured mechanism that ensures faithful post-mitotic NPC assembly and that does not depend on a sole nucleoporin. At a closer look, however, Mad1- and Nup153-depleted cells showed prominent ultrastructural changes in membrane curvature at the NPC-NE interface (Fig. 4A,B) and in NE spacing (Fig. 4A-D). Indeed, the yeast ortholog of Nup153, Nup1, is required for correct membrane curvature due to an N-terminal amphipathic helix (Meszaros et al., 2015) and Nup153 similarly comprises such an amphipathic helix in its N terminus (Vollmer et al., 2015). By their amphipathic helices both nucleoporins are capable of directly interacting with liposomes *in vitro* and inducing liposome tubulation (Meszaros et al., 2015; Vollmer et al., 2015), suggesting an evolutionary conserved necessity for Nup153’s amphipathic helix for correct membrane curvature at NPCs. Similarly, other nucleoporins containing amphipathic helices, such as Nup155, Nup133, and Nup53 are capable of inducing membrane curvature (Drin et al., 2007; Kim et al., 2014; Schwartz, 2016; Vollmer et al., 2012). The expansion of the spacing between outer nuclear membrane (ONM) and inner nuclear membrane (INM) is likely caused by an increase in the tension between lipids in the NE, which is normally attenuated by amphipathic helices, such as the one in Nup153. Implicated in NE spacing have been also proteins forming LINC complexes to connect the cytoplasm to the nucleoplasm: SUN (Sad1 and UNC-84) proteins in the INM and KASH (Klarsicht, ANC-1, and Syne homology) proteins in the ONM (Crisp et al., 2006). SUN proteins, however, have been shown to not dictate the width of the NE and SUN-KASH bridges are only required to maintain NE spacing in cells subjected to increased mechanical forces (Cain et al., 2014). Consistently, we observed a regular localisation of both SUN and KASH proteins in the NE of Nup153- and Mad1-depleted cells (Fig. S4B).

Do these membrane curvature and NE spacing abnormalities reflect post-mitotic or interphase assembly defects? We consider it likely as post-mitotic defects, as defects in interphase assembly may rather lead to membrane collapse than expansion, as describe for POM121 (Talamas and Hetzer, 2011). Furthermore, the depletion of Nup153 led to a displacement of the cytoplasmic nucleoporins Nup358, Nup214, and Nup88 (Figs 5 and 6) as well as to reduced NPC density after S phase (Fig. 7). Recruitment of cytoplasmic nucleoporins to NPCs was also seen in Nup1-depleted yeast cells (Meszaros et al., 2015). NPC assembly in yeast occurs exclusively into a closed NE, strongly suggesting that the displacement of the cytoplasmic nucleoporins in cells lacking Nup153 is due to interphase NPC assembly defects. It is conceivable that newly synthesised cytoplasmic nucleoporins cannot incorporate into NPCs during the G2 phase of the cell cycle, when the NE is expanding after DNA replication (Makio et al., 2009; Meszaros et al., 2015; Vollmer et al., 2015). Displacement of cytoplasmic nucleoporins and a reduction in NPC density was not observed in Mad1-depleted cells, suggesting that Nup153 acts independently of Mad1 in interphase NPC assembly. Together our data hence suggest that Nup153 has separable roles in post-mitotic NE formation in concert with Mad1 and in interphase NPC assembly independently of Mad1. Whether Mad1’s role in post-mitotic NE reformation is solely to its role in Nup153 recruitment to anaphase chromatin remains to be elucidated.

## Material and Methods

All experiments were carried out at room temperature, unless otherwise stated.

### Cell culture and transfections

HeLa cells and 3T3 cells were grown in Dulbecco’s modified Eagle’s medium (DMEM) supplemented with 10% foetal bovine serum (FBS; Biochrom GmbH, Berlin, Germany), 100 U/ml penicillin (Thermo Fisher Scientific, Invitrogen, Waltham, MA, USA) and 100 U/ml streptomycin (Invitrogen) and maintained at 37°C in a humidified incubator with 5% CO_2_. HeLa cells stably expressing YFP-tubulin were grown in DMEM containing 10% FBS, penicillin, streptomycin and 250 μg/ml geneticin (G418).

HeLa T-Rex cells expressing GFP-Nup98 were established by transfection with pcDNA4/TO-GFP-NUP98 and positive clones were selected by treatment with 5 mg/ml blasticidin and 200 mg/ml zeocin. Individual clones were isolated, expanded and cultured in MEM (Life Technologies Gibco, Gent, Belgium) containing 10% FBS, blasticidin, zeocin, penicillin and streptomycin. HeLa T-Rex cells expressing GFP-Nup153 (Duheron et al., 2014) or GFP-Nup98 were grown in Minimal Essential Medium (MEM) containing 10% FBS, penicillin, streptomycin, 5 μg/ml blasticidin (Invitrogen) and 200 μg/ml zeocin (Invivogen, Toulouse, France). Cells were treated with 1 μg/ml of tetracycline (Sigma-Aldrich, Machelen, Belgium) for 24 h to induce the expression of GFP-Nup153 or GFP-Nup98.

Cells were transfected using Lipofectamine RNAiMAX (Invitrogen) for siRNAs and Lipofectamine 2000 (Invitrogen) for plasmids according to the instructions of the manufacturer. The siRNAs used were On-target plus smart pool siRNAs purchased from Dharmacon (Lafayette, CO, USA): Non-targeting siRNAs (D-001810-10), human Nup153 siRNAs (L-005283-00), human Tpr siRNAs (L-010548-00), human Mad1 siRNAs (L-006825-00), human Mad2 siRNAs (L-003271-00), mouse cyclophilin B control (D-001820-02), mouse Nup153 siRNAs (L-057025-01).

To arrest cells at the G2/M boundary, cells were treated with 9 μM RO-3306 (Sigma-Aldrich) for 18 h.

### Constructs

For the generation of pGEX-6P-Nup153, human *NUP153* was amplified by PCR and inserted into XhoI/NotI cut pGEX-6P. For the generation of pEGFP-Nup153, PCR-amplified human *NUP153* was inserted into HindIII/XmaI cut pEGFP-C3. For all Mad1 constructs, human *MAD1* was amplified by PCR. Recombinant full-length FLAG-Mad1 and FLAG-Mad1-N596 were subcloned from pFLAG-CMV2-Mad1 into NheI/BamHI cut pET24d+. The other fragments were generated by adding an N-terminal FLAG-tag into the forward primers. The resulting products were subcloned into NheI/BamHI cut pET24d+. All the constructs were verified by DNA sequencing. All primers are listed in Table S1.

### Antibodies

Primary antibodies used were: monoclonal mouse anti-Nup153, clone SA1 (hybridoma supernatant, a kind gift from Dr. Brian Burke, University of Singapore; IF 1:100, WB1:50), mouse anti-Mad1 (Santa Cruz, Heidelberg, Germany, sc-47746; IF 1:100, WB 1:1000;), mouse anti-Tpr (Abnova, Hamm, Germany, H00007175-M01; IF 1:400, WB 1:2000), mouse anti-Mad2 (Sigma-Aldrich, M8694; WB 1:1000), mouse anti-laminA/C (Abcam, Cambridge, UK, ab40567; IF 1:60), mouse anti-FLAG (Sigma-Aldrich, F3165; WB 1:2000), mouse anti-Hec1 (Abcam, ab3613; IF 1:200), mouse mAb414 (Covance, Mechelen, Belgium; IF 1:5000); polyclonal rabbit anti-Nup153 (Sigma-Aldrich, HPA027896; IF 1:400), rabbit anti-Mad1 (Santa Cruz, sc-67338; IF 1:100), rabbit anti-Mad2 (Covance, PRB-452C; IF 1:200), rabbit anti-α-tubulin (Abcam, ab18251; WB 1:4000), rabbit anti-actin (Sigma-Aldrich, A2066; WB 1:1000), rabbit anti-Nup88 (BD Biosciences, Erembodegem, Belgium, 611896; IF 1:500), rabbit anti-Nesprin 2 (a kind gift of Dr. Iakowos Karakesisoglou; IF 1:300) rabbit anti-Sun1 (a kind gift of Dr. Ulrike Kutay; IF 1:1000), rabbit anti-Nup214 (a kind gift of Dr. Ralph Kehlenbach; IF 1:1000), rabbit anti-Nup358 (a kind gift of Dr. Mary Dasso; IF 1:300).

Secondary antibodies for immunofluorescence were: goat anti-mouse IgG-Alexa 488, goat anti-rabbit IgG-Alexa 488, goat anti-mouse IgG-Alexa 568, goat anti-rabbit IgG-Alexa 568, chicken anti-mouse IgG-Alexa 647 from Molecular Probes (Paisley, UK). All antibodies were used at a dilution of 1:1000. For Western blot, secondary goat anti-mouse IgG and goat anti-rabbit coupled with alkaline phosphatase antibodies (Sigma-Aldrich) were used at a dilution of 1:20.000.

### GST pull-down assays

All recombinant FLAG-Mad1 fragments, GST, and GST-Nup153-N were produced in *E. coli* BL21 codon plus (DE3) cells. Bacteria precultures were grown overnight at 37°C in 1 ml LB medium containing appropriate antibiotics and diluted into 100 ml LB medium containing appropriate antibiotics. Protein expression was induced with 0.1 mM of isopropyl-beta-D-thiogalactopyranoside (IPTG) at an optical density at 600 nm of 0.5 and cells were grown for further 3 h at 37°C. The cells were collected by centrifugation at 4°C at 3220 g for 20 min, resuspended in 4 ml PBS containing 1% Triton X-100 plus protease inhibitor cocktail and next sonicated on ice (5 times 10 sec, with 10 sec off between each sonication). After centrifugation at 4°C at 16.000 g for 15 min, the supernatants were collected, frozen in liquid nitrogen, and stored at −80°C.

For pull-down assays, 500 μl of GST or 200 μl of GST-Nup153-N were bound to 20 μl of glutathione-sepharose beads for 1 h at 4°C on a rocker platform. Next, the beads were washed twice with PBS containing 1% Triton X-100 and a cocktail of protease inhibitors and subsequently twice with 50 mM HEPES, pH 7.4, 150 mM NaCl, 1 mM DTT, 0.1% NP-40 and protease inhibitors. The beads were next incubated with the different recombinant FLAG-Mad1 fragments for 1 h at 4°C on a rocker platform. After binding, beads were washed four times with 50 mM HEPES, pH 7.4, 150 mM NaCl, 1 mM DTT, 0.1% NP-40, resuspended in 2x Laemmli buffer (125 mM Tris-HCl, pH 6.8, 4% SDS, 20% glycerol, 10% 2-mercaptoethanol, 0.004% bromophenol blue) and boiled for 5 min at 95°C. Next, the samples were separated by SDS-PAGE and analysed by Western blot.

### Immunofluorescence microscopy

Cells were grown on glass coverslips and fixed with 2% formaldehyde for 15 min. Next, cells were washed three times with PBS for 5 min and permeabilised with PBS containing 0.2% Triton X-100 and 1% bovine serum albumin (BSA) for 10 min. After three washes with PBS containing 1% BSA for 5 min, cells were stained with the appropriate antibodies for 2 h at room temperature or overnight at 4°C in a humid chamber. Next, cells were washed three times with PBS/1% BSA, incubated with secondary antibodies for 1 h and washed four times with PBS. The coverslips were then mounted with Mowiol-4088 (Sigma-Aldrich) containing 1 μg/ml DAPI and stored at 4°C until viewed. Images were acquired using a 63x oil immersion objective on a LSM710 laser-scanning confocal microscope (Zeiss, Oberkochen, Germany) or on an Axio Observer Z.1 microscope (Zeiss). Images were recorded with the respective system software and processed using Image J and Adobe Photoshop.

To visualise importin ß at the NE, cells were washed once with PBS and then with ice-cold buffer (20 mM HEPES, pH 7.3, 110 mM potassium acetate, 2mM magnesium acetate). Next, cells were permeabilized using freshly made ice-cold buffer complemented with 2 mM DTT, protease inhibitors cocktail and digitonin at a concentration of 40 μg/ml. After 5 min incubation on ice, cells were washed twice with PBS and subjected to immunofluorescence as described above.

### Proximity ligation assays

Proximity ligation assays were performed according to the instructions of the manufacturer (Duolink; Olink Bioscience, Uppsala, Sweden). HeLa cells were grown on glass coverslips, fixed, and permeabilized as described above for immunofluorescence experiments. After incubation with primary antibodies, the cells were subjected to the Duolink red kit. After washing with Duolink wash buffer A, cells were incubated with the Duolink PLA probes for 1 h at 37°C in a pre-heated humidified chamber. Following washing steps with Duolink wash buffer A, the Duolink ligation reagent was incubated for 30 min followed by the Duolink amplification reagent for 90 min, both at 37°C in a pre-heated humidified chamber. Cells were washed with Duolink wash buffer B and mounted with the Duolink *in situ* mounting medium containing DAPI. Images were acquired using a 63x oil immersion objective on a Zeiss Axio Observer Z.1 fluorescence microscope, recorded with Axiovison software and processed using ImageJ and Adobe Photoshop.

### Time-lapse imaging

HeLa T-Rex were grown in CELLview cell culture chambers with glass bottom (Greiner Bio-One, Kremsmünster, Austria), transiently treated with siRNAs for 48 h, and maintained at 37°C in complete MEM medium. Next, to induce expression of GFP-Nup153 or GFP-Nup98, cells were treated with tetracycline at a final concentration of 1 μg/ml for 24 h. Cells were equilibrated in a CO_2_-independent medium without phenol red (Leibovitz’s L-15 medium; Gibco) supplemented with 10% FBS, penicillin and streptomycin, and L-glutamine. About 2 h after medium exchange, cells were placed into a 37°C pre-heated incubation chamber. Time-lapse sequences were recorded every min during around 1 h. Images were taken using a 63x/1.4 oil immersion objective lens on a LSM710 laser-scanning confocal microscope (Zeiss) and were recorded using the system software. The images were processed with ImageJ and Adobe Photoshop.

### Super-Resolution 3D Structured Illumination microscopy

HeLa cells were grown on glass coverslips and treated with siRNAs for 48 h and subjected to immunofluorescence as described above. The coverslips were mounted using Vectashield 1000 and sealed with nail polish. For acquisition, the 488 and 568 laser lines were used. Images were taken on DeltaVision OMX-Blaze version4 microscope (Applied Precision, Issaquah, WA). The images were processed and reconstructed with the DeltaVision OMX SoftWoRx software package (Applied Precision).

### Electron microscopy

HeLa T-Rex cells were grown in 6-well plates and treated with the respective siRNAs and maintained in 5% CO_2_ at 37°C in MEM. 48 h after transfection (72 h for Tpr siRNAs), cells were collected using a cell scraper, pelleted, and washed once with PBS. Cells were fixed for 1 h in Karnofski solution (10 mM PBS, pH 7.4, 3% paraformaldehyde, 0.5% glutaraldehyde) and washed once with PBS. Post-fixation was performed in 1% reduced osmium tetroxide containing 1.5% potassium ferricyanide for 40 min and 1% osmium tetroxide for another 40 min. Cells were washed with water and samples were next dehydrated in a series of ethanol solutions, embedded in Epon resin, and subjected to EM analysis. EM micrographs were taken on a Phillips CM-100 transmission electron microscope, equipped with a CCD camera, at an acceleration voltage of 80 kV. Images were recorded using the system software and processed using Adobe Photoshop.

### Western blots

Cells were harvested by trypsinisation, washed with PBS, and resuspended in lysis buffer (50 mM Tris-HCl, pH 7.5, 150 mM NaCl, 1% NP-40 supplemented with a cocktail of protease inhibitor tablets (Roche, Basel, Switzerland)) and incubated for 15 min at 4°C. Subsequently, the lysates were cleared by centrifugation at 16.000 g at 4°C for 10 min. The lysates were resuspended in Laemmli buffer and boiled for 5 min at 95°C. 20 μg of proteins were then separated by SDS-PAGE for 90 min at 100 V and the proteins were transferred onto a PVDF membrane with a current of 20 mA overnight. Next, the membrane was incubated for 1 h in TBS containing 0.1% Tween 20 (TBS-T) and 5% of non-fat dry milk, followed by incubation of primary antibodies in the blocking solution for 2 h at room temperature or overnight at 4°C. After washing three times with TBS-T, the membrane was incubated with the appropriate alkaline phosphatase-conjugated secondary antibody for 1 h. After three washes with TBS-T, the membrane was washed twice with assay buffer (100 mM Tris-HCl, pH 9.8, 10 mM MgCl_2_) for 2 min. The membrane was incubated for 5 min with the Lightning CDP Star Chemiluminescence reagent (ThermoFisher Scientific, Applied Biosystem) and exposed on CL-Xposure film (ThermoFisher Scientific).

### Nocodazole arrest and spindle assembly checkpoint analysis

Cells were grown in 6-well plates and transiently treated with siRNAs. 24 h post transfection, cells were treated with 2 mM thymidine (Sigma-Aldrich) for 24 h, washed with PBS, released into fresh medium for 3 h, and treated with 100 ng/ml nocodazole (Sigma-Aldrich) for 12 h. Cells were next released into fresh medium for 0, 45, 90, 120, and 150 min, respectively, lysed in lysis buffer (50 mM Tris-HCl, pH 7.5, 150 mM NaCl, 1% NP-40 supplemented with a cocktail of protease inhibitor tablets (Roche)) and subjected to Western blot analysis.

## Acknowledgements

The authors thank Drs. Brian Burke (A* STAR Biomedical Sciences Cluster, Singapore), Vincent Duheron (Université Libre de Bruxelles, Belgium), Iakowos Karakesisoglou (University of Durham, UK), Ralph Kehlenbach (University of Göttingen, Germany), and Ulrike Kutay (ETH Zurich, Switzerland) for sharing reagents. We are grateful to Dr. Alexia Loynton-Ferrand from the Imaging Core Facility and Ursula Sauder and Vesna Oliveri from the BioEM Facility of the Biozentrum, University of Basel, Switzerland, for SIM and electron microscopy expert technical assistance. Confocal images were acquired at the CMMI (Charleroi, Belgium), which is supported by the European Regional Development Fund (ERDF).

This work was supported by a FRIA PhD fellowship from the Fonds de la Recherche Scientifique-FNRS Belgium to IM and by research grants to BF (grant numbers F.6006.10 and T.0237.13).

